# aBayesQR: A Bayesian method for reconstruction of viral populations characterized by low diversity

**DOI:** 10.1101/103630

**Authors:** Soyeon Ahn, Haris Vikalo

**Affiliations:** The University of Texas at Austin, Austin TX, USA

**Keywords:** viral quasispecies reconstruction, low diversity, bayesian method

## Abstract

RNA viruses replicate with high mutation rates, creating closely related viral populations. The heterogeneous virus populations, referred to as viral quasispecies, rapidly adapt to environmental changes thus adversely affecting efficiency of antiviral drugs and vaccines. Therefore, studying the underlying genetic heterogeneity of viral populations plays a significant role in the development of effective therapeutic treatments. Recent high-throughput sequencing technologies have provided invaluable opportunity for uncovering the structure of quasispecies populations (i.e., reconstruction of viral sequences and discovery of their relative frequencies). However, accurate reconstruction of viral quasispecies remains difficult due to limited read-lengths and presence of sequencing errors. The problem is particularly challenging when the strains in a population are highly similar, i.e., the sequences are characterized by low mutual genetic distances, and further exacerbated if some of those strains are relatively rare; this is the setting where state-of-the-art methods struggle. In this paper, we present a novel viral quasispecies reconstruction algorithm, aBayesQR, that employs a maximum-likelihood framework to infer individual sequences in a mixture from high-throughput sequencing data. The search for the most likely quasispecies is conducted on long contigs that our method constructs from the set of short reads via agglomerative hierarchical clustering; operating on contigs rather than short reads enables identification of close strains in a population and provides computational tractability of the Bayesian method. Results on both simulated and real HIV-1 data demonstrate that the proposed algorithm generally outperforms state-of-the-art methods; aBayesQR particularly stands out when reconstructing a set of closely related viral strains (e.g., quasispecies characterized by low diversity).

## 1 Introduction

A number of potentially life-threatening infectious diseases are caused by RNA viruses, including human immunodeficiency virus (HIV), hepatitis C virus (HCV), influenza and Ebola. RNA viruses have a relatively high mutation rate due to both their error-prone replication process and the lack of sophisticated repair mechanisms [1]. Consequently, they rapidly evolve and exist as a set of non-identical but closely related genetic variants, known as a viral quasispecies. Viral populations can readily adapt to dynamic environments and develop resistance to antiviral drugs and vaccines, which makes the design of effective and long-lasting treatments for RNA viral diseases exceedingly difficult [2]. Determining the structure of viral populations helps the understanding of viral diseases and provides guidance in the development of effective medical therapeutics. Quasispecies spectrum reconstruction (QSR) aims to assemble individual haplotype sequences in a population and estimate their prevalence using sequencing reads generated from a sample containing a set of viral variants. High-throughput next-generation sequencing (NGS) technologies have enabled affordable acquisition of data needed to assemble quasispecies. However, relatively short length of the NGS reads and the presence of errors in sequencing data render the QSR problem difficult. The QSR problem is particularly challenging when the strains in a viral population are highly similar, i.e., the sequences are characterized by low mutual genetic distances, and further exacerbated if some of those strains are relatively rare [3].

Several software tools for solving the QSR problem by analyzing NGS data have been developed in recent years. ShoRAH [4], the earliest publicly available such software, was developed by combining a path cover based approach and probabilistic clustering in [5] and [6], respectively, and applied to analysis of HIV data [7]. Read-graph approach was the basis for ViSpA [8], developed as a variant of the network flow method proposed in [9]. [10], proposed a combinatorial method for QSR and the resulting software, QuRe, was provided by [11]. An approach that resulted in the software package PredictHaplo [12] relied on a Dirichlet Process mixture model and was developed specifically targeting HIV population reconstruction; QuasiRecomb [13] is based on a hidden Markov model that explicitly models recombination events. In [14], a benchmarking study that compares the performance of several publicly available quasispecies reconstruction softwares was presented. The study demonstrated that none of the tested methods could reconstruct populations characterized by low pairwise distance between the haplotype sequences. Following this study other softwares, including HaploClique [15], based on max-clique enumeration of a read alignment graph, and VGA [16], a graph-coloring based heuristic method, were developed. Most recently, a reference-assisted *de novo* assembly pipeline, ViQuaS, was proposed in [17]. ViQuaS extends an existing algorithm, QuRe [10], and outperforms various other techniques on a wide range of dataset. However, performance of these more recent methods deteriorates dramatically in the scenarios where the genetic diversity of a population is low [3].

Both [3] and [14] have pointed out that the existing methods for viral quasispecies reconstruction struggle in the scenarios where the populations are characterized by low diversity. This, in part, is due to the presence of relatively long genetic regions that are common to pairs of closely related viral sequences; clearly, this makes distinguishing different strains challenging. The problem becomes even more difficult when the frequency of one (or more) of the close strains is low; in such settings small genetic distances may be confused for sequencing errors and hence remain undetected. Such failures to detect may have serious consequences in antiviral treatment studies since undetected strains cannot be properly targeted for drug and vaccine design. It has been shown that even the viral strains existing at low frequencies can cause a drug treatment failure due to their resistance to the drug [18, 19]. Therefore, complete recovery of the composition of viral populations is of critical importance for effective antiviral therapies.

In this paper, we propose a novel QSR algorithm, aBayesQR (combining agglomerative hierarchical clustering and Bayesian inference), that overcomes limitations of the existing methods and reliably reconstructs quasispecies characterized by low diversity. The algorithm performs reconstruction of a quasispecies from next-generation sequencing (NGS) data in two stages. In the first stage, conflict-free short reads are hierarchically merged and assembled into longer sequences (contigs) which we refer to as super-reads. In the second stage, likelihoods of the probable quasispecies are computed using the assembled super-reads (rather than using the original set of short reads), and the most likely set of viral strains is selected. Note that the super-reads synthesized in the first stage of aBayesQR allow us to distinguish between closely related strains which share long genetic regions as well as reduce the search space and enable computational tractability of the Bayesian inference conducted in the second stage. The second stage of aBayesQR involves sequential pruning of the solution space; in particular, the likely set of partial viral strains comprising *n* single nucleotide variants (SNVs) is generated by extending previously inferred partial viral strains having *n* – 1 SNVs. The number of sequences in a set (i.e., the size of a viral population) is dynamically updated at each step by evaluating quality of the set of partially reconstructed viral strains, and ultimately precisely inferred at the end of the search process. The relative frequencies of each strain are determined by counting the numbers of reads unambiguously associated with each of the reconstructed strains. Our tests on both simulated and experimental data demonstrate superior performance compared to state-of-the-art methods for quasispecies reconstruction. In particular, it is shown that unlike the competing methods, aBayesQR is capable of detecting and reliably reconstructing viral haplotypes having very small mutual genetic distances.

## 2 Proposed Method

Our algorithm for inferring spectrum of a viral population consists of the following two steps: (1) constructing super-reads by hierarchically clustering aligned paired-end reads, (2) inferring the most likely quasispecies from the set of superreads and estimating the frequencies of the strains in the quasispecies.

### 2.1 Super-reads construction via agglomerative clustering

In the first stage of aBayesQR, paired-end reads uniquely mapped to a reference genome are grouped into super-reads via agglomerative hierarchical clustering. This is facilitated by a weighted graph 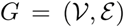 which is constructed and recursively updated as the clustering proceeds. In particular, each vertex of *G* is associated with a cluster collecting reads that originated from a single strain in a quasispecies; we denote the set of reads in the *i^th^* cluster (i.e., the cluster associated with the *i^th^* vertex) as 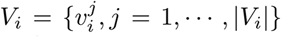. Let *sr_i_* denote a consensus sequence (i.e., a super-read) constructed from the reads in *V_i_*. The *i^th^* and *j^th^* vertex of *G* are connected by an edge *e_ij_ ∈ ε* if all the reads in *V_i_* and *V_j_* (or, equivalently, *sr_i_* and *sr_j_*) are conflict-free and an overlap criterion, specified later in this subsection, is satisfied. The weight *w_ij_*of the edge *e_ij_* is a measure of similarity between *V_i_* and *V_j_* at each step, the algorithm merges a pair of vertices connected by the edge having the largest weight to form a new vertex and agglomerates the corresponding clusters.

The alleles at homozygous sites, common to all the components of a quasispecies, are not utilized in the reconstruction procedure. Instead, we separate reads having originated from different strains by clustering them using heterogeneous sites with reliable SNV information. An SNV information is considered reliable if the relative abundance of the allele is above a pre-determined threshold, as in ([20]); alleles whose abundance is below the threshold are treated as sequencing errors and disregarded in the process of clustering. For convenience, let us denote the set of pre-processed paired-end reads by *R* = {*r_i_, i* = 1,…, |*R*|}. The agglomerative clustering is initialized with *|R|* clusters, one for each read; in other words, we start with *V*_1_ = *r*_1_,…, *V*_|*R*|_= *r_|R|_*, implying that 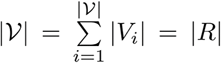, and proceed by sequentially merging judiciously chosen pairs of vertices (i.e., agglomerating the corresponding clusters). Intuitively, it is meaningful to reduce the number of vertices in the graph by merging those associated with conflict-free consensus sequences that have a large overlap. To formalize this, let 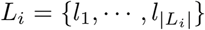 denote an index set of the SNV positions covered by *sr_i_*, let 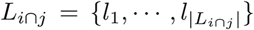 covered by both *sr_i_* and *sr_j_*, and let 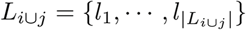 be the index set of SNV positions be the index set of SNV positions covered by either *sr_i_* or *sr_j_*. Then the pairs of vertices (*i, j*) that we consider as candidates for merging and thus connect by an edge are those satisfying either

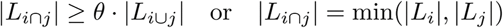

where the 2*^nd^* condition promotes merger of short super-reads, and the choice of *θ* is discussed below. To quantify uncertainty inherent to a clustering solution due to existence of non-overlapping positions among the reads in each cluster, we define a position-specific confidence score

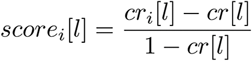

where *l* denotes the position, *cr*[*·*] is the overall coverage rate, and *cr_i_*[*·*] denotes cluster-specific coverage rate for *V_i_* (i.e., *cr_i_*[*l*] is the fraction of reads in 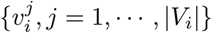 covering position *l*). On the one hand, this score is penalized at a site where the fraction of cluster members (short reads) covering the site is low; the score is negative if the cluster-specific coverage rate is below the global coverage rate which implies uncertainty of the clustering decision. On the other hand, positive scores indicate high confidence in the decision to group the reads into the same cluster. Note that the highest possible score of 1 at position *l* is achieved when all the reads in a cluster cover the *l^th^* position. Using the confidence scores, we define the weight *w_ij_* assigned to an edge *e_ij_* to quantify similarity between *V_i_* and *V_j_* as

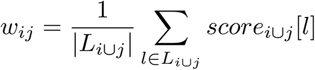

Given the weights *w_ij_*, we can now specify the clustering procedure. In each step, the pair of vertices connected by the edge with maximum weight is merged; the newly constructed vertex inherits edges from the merged vertices and the weights on those edges are re-evaluated. A new (longer) consensus sequence is constructed by combining the two super-reads associated with the merged vertices; recall that there are no conflicts between the super-reads being merged. If after such an update step no edges connect the new vertex with the rest of the graph (because no inherited edges satisfy the connectivity condition), *θ* is decreased and the above process is repeated. We initially set *θ* to 0.9 and gradually decrease it by 0.1 while *θ >* 0. The above procedure is repeated until no pairs of vertices satisfy the connectivity condition. By that point, a set of long consensus sequences (the final super-reads) has been formed from the clusters of reads associated with the nodes of the final graph. While the complexity of agglomerative clustering is, in general, *O*(*N* ^3^) where *N* denotes the input data size ([21]), it has been shown that its time complexity can be reduced to *O*(*N* ^2^) with accuracy equal to that of the brute-force method by using the partial maximum array technique [22]. We exploit this to efficiently construct super-reads. The algorithm for super-read construction is formalized as Algorithm 1.

### 2.2 ML reconstruction of quasispecies from super-reads

Here we describe how to reconstruct the most likely set of strains in a viral quasispecies using super-reads from Sect. 2.1 and their confidence scores. While in principle the method outlined in this section could be applied directly to the short reads provided by a sequencing platform, such an approach would in general not only be computationally prohibitive due to a very large number of short reads but also limit the ability of the algorithm to distinguish strains with small mutual genetic distances due to having long conserved regions. Relying on a relatively small number of long super-reads constructed from short reads circumvents both of these problems and makes the reconstruction more accurate and practically feasible. Note that sequencing errors may undesirably prevent clusters of reads from being merged with other clusters due to a violation of conflict-free requirement; consequently, a set of short reads in a small cluster is likely to have a disproportionate amount of sequencing errors. For this reason, we ignore clusters with very small memberships (in particular, those containing fewer than 0.001 · |*R*| reads), which limits the detection of strains to those constituting more than 0.1% of the quasispecies.

#### Algorithm 1 Agglomerative clustering for super-reads construction

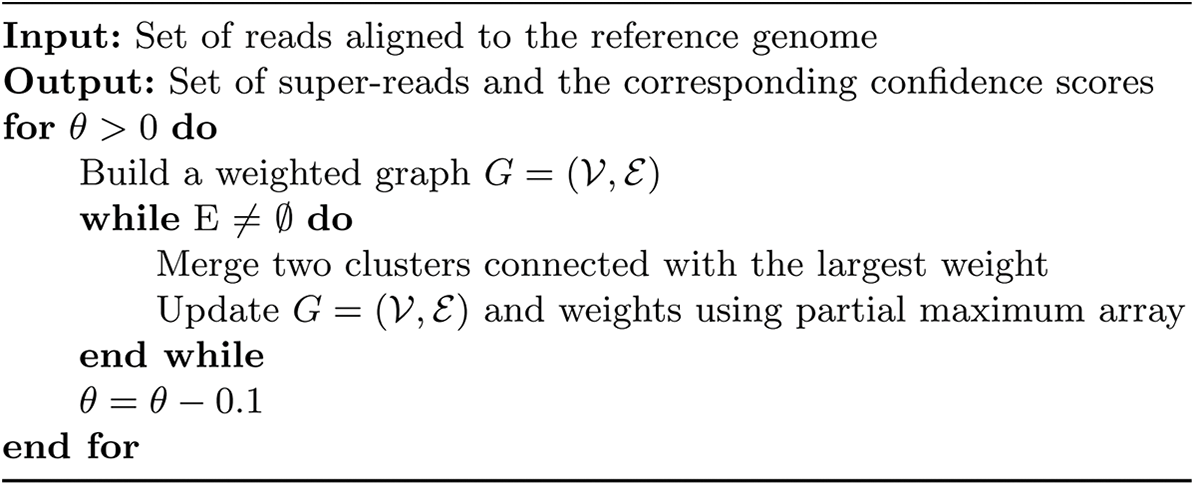

Let *C* = {*C_m_, m* = 1,…,*M*} denote the collection of clusters that remain after deleting clusters having only few reads; moreover, for convenience let us re-label the reads in *C_m_* as 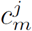, i.e, 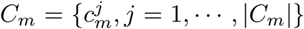 where 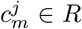. We organize the super-reads obtained by Algorithm 1 in Sect. 2.1 into the rows of an *M* × *N* matrix **S** = {*s_mn_, m* = 1,…, *M*, *n* = 1,…, *N*} with entries *s_mn_ ∈* {A, C, G, T, –} where – denotes a site not covered by a super-read and *N* denotes the total number of SNV sites in the strains of a quasispecies. A nucleotides in the (*m, n*) position of **S** is assigned confidence *score_m_*[*n*] defined in Sect. 2.1; the scores for the entire matrix are normalized so that they fall between 0 and 1 in order to use them in our Bayesian approach to assembly. Let *ε_mn_* be the probability that *s_mn_* was estimated erroneously due to either a sequencing error in reads on the *n^th^* SNV position or the uncertainty induced by reads not covering the *n^th^* SNV position. Note that negative scores indicates low confidence resulting from insufficient cluster-specific coverage rate while positive scores imply relatively confident information. In order to map *score_m_*[*n*] *∈* (*−∞,* 1] to the set [0, 1], we set 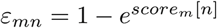 for *score*_m_[*n*] < *ln*(1 – *ε*), where *ε* denotes the error rate of a sequencing platform. Otherwise, we set *ε_mn_*=*ε*.

Let *Q* = {*q_k_*, *k* = 1,…, *K*} denote the set of *K* strains of a viral quasispecies. The goal in the second stage of our method is to determine *Q* from the super-reads matrix **S** using a probabilistic framework. An exhaustive search over the entire solution space is computationally intractable even for small **S**; instead, we reconstruct the set of *K* viral strains sequentially, extending partially estimated strains one SNV position at each step. Since maintaining and extending all possible partial strains inevitably increases their number exponentially, unlikely sets of candidate strains are pruned in each step. Each step consists of three basic parts: (*a*) extension of the partially reconstructed strains, (*b*) selection of probable sets comprising *K* strains chosen among those generated in step (a), and (*c*) evaluation of the quality of the selected sets of strains and an update of *K*. The sequential Bayesian inference procedure in step *t* is illustrated in Fig. S1 in Appendix A.

### Extending partially reconstructed strains

Let 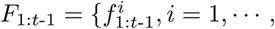|*F*_1:*t*-1_| be the collection of partially reconstructed strains covering the first *t −* 1 SNV sites and let 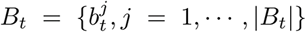 be the lists of distinct bases in the *t^th^* column of **S**, where 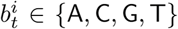 and 2 ≤ |B*_t_*| ≤ 4. Then, all the possible extensions of 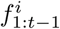 to the SNV site *t* can be enumerated as 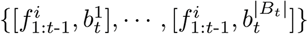. Let 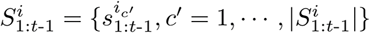 be the collection of super-reads covering some of the first *t* SNV sites which are consistent with 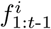 (ignoring “*−*” in 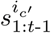) where {*i_c´_*} denote indices of rows of **S** that are placed in 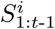 and let 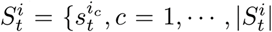} denote the collection of nucleotides ( 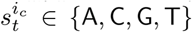, not “*−*”) observed at the *t^th^* SNV site of the super-reads in 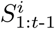 where {*i_c_*} denote the indices of rows in **S** that contribute to 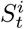. Given 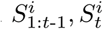 and 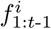, the probability of 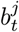 being the true extension of 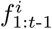 is given by

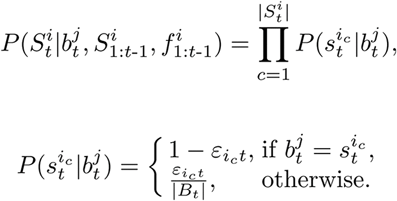

We extend 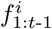 to 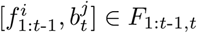 by appending the 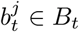 which satisfies 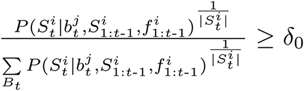, where the exponent ensures proper normalization and is needed since the number of super-reads, 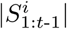, varies for each 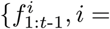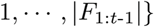. For 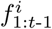 which has no matched super-reads, i.e., 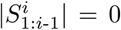, we keep all of |*B_t_*| possible extensions of 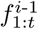. By collecting probable extensions for each 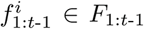, we obtain the set of partial strains stretching over the first *t* SNV sites, 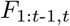. This procedure is formalized as function *ExtendFrag* in Appendix A.

### Inferring likely sets of K partial strains

Having generated the probable partial strains *F*_1:*t*-1,*t*_, we denote the set of all its possible subsets of *K* strains (i.e., the quasispecies population candidates) as 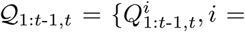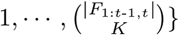 where 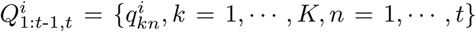 and 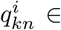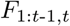. The log-likelihoods of 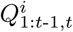 can be expressed as

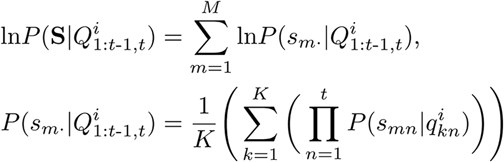

where *s_m_*. denotes the *m^th^* row vector of the matrix of super-reads **S** and

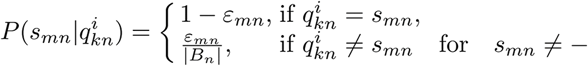

Let 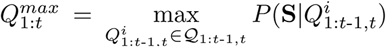. Among the 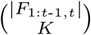 sets in 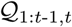, we keep only those that satisfy 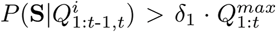 while the others are discarded; let us denote the collection of candidate sets that pass this test as 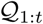. For practical feasibility of the scheme, the collection of partially reconstructed strains *F*_1:*t*-1,*t*_ is trimmed by excluding from it all the strains that are not part of at least one of the sets in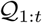; we denote the resulting collection of partial strains by *F*_1:*t*_ ∈ *F*_1:*t*-1,*t*_ and use it when extending the strains onto the *t* + 1 SNV site. The described procedure is formalized as function *InferQuasi* in Appendix A.

### Determining the number of strains *K* in a quasispecies

In this step, we assess appropriateness of *K* used in the inference of 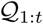 and update it if necessary. To this end, we rely on the minimum error correction (MEC) score which has previously been broadly used as a criterion in the design of methods for haplotype assembly ([23] and [24]). In the context of polyploid haplotype assembly, the MEC score is defined as the smallest number of nucleotides that needs to be changed in data (i.e., in observed reads) so that the corrected reads are consistent with having originated from *K* haplotypes. Let *HD_t_*( ·, ·) denote the Hamming distance between two sequences counted over the observed nucleotides in the first *t* SNV positions.^1^ Then the MEC score of the most likely set 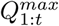 of *K* viral strains evaluated on the first *t* SNVs is

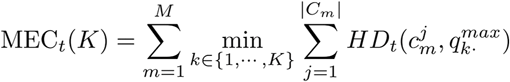

where 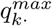 is the *k^th^* row vector of 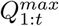. Let *N_t_* be the total number of nucleotides observed in the first *t* SNV positions of all the reads of the dataset.

Note that the smaller the MEC scores, the higher the accuracy of a clustering. If MEC*_t_* (*K*)*/N_t_ <* 2є, we use the same value *K* in the next step where the likely set of viral strains stretching over the first *t* + 1 SNV positions is inferred. Otherwise, we increase *K* by 1, repeat the estimation of 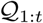, and evaluate the improvement rate of MEC score as

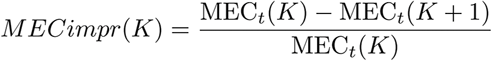

The reason for selecting *K* based on the MEC improvement rate (*M ECimpr*) is that the MEC score drops significantly once *K* matches the actual number of clusters; our scheme attempts to detect that change in order to infer population size. If *M ECimpr* (*K*) *> η*, where *η* denotes a pre-specified threshold, the number of species is updated as *K* ← min{*K* + *n*, |*F*_1:*t*-1,*t*_|}where *n* is the smallest integer number such that *MECimpr* (*K* + *n*) *< η*. If *MECimpr* (*K*) *< η*, we update the number of species as *K* ← max*{K* − *n,* 2*}* where *n* is the smallest integer such that *MECimpr* (*K − n*) *≥ η*. The choice of threshold *η* is discussed in the Appendix B. The updated value of *K* is used for the inference of 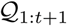. Note that the probable set of viral strains, 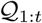, is stored for each *K* to avoid performing redundant *M ECimpr*(·) calculations.

Once we obtain the most likely set of *K* viral sequences covering *N* SNVs, 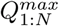, the full-length *K* quasispecies strains are reconstructed by inserting the consensus nucleotides observed in *R* into the non-SNV sites. We estimate relative frequencies *p_k_*, 1 ≤ *k* ≤ *K*, of quasispecies strains based on the Hamming distance between super-reads and the reconstructed sequences. In particular, for each super-read *sr_i_* we determine the nearest assembled strain *q_j_* where 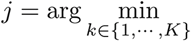*HD*(*sr_i_, q_k_*) and the number of reads involved in constructing the super-read *sr_i_* is counted towards *p_j_*. The entire scheme proposed in this subsection is summarized as Algorithm 2.

## 3. Results and Discussion

### 3.1 Performance comparison on simulated data

To evaluate performance of the proposed method for quasispecies reconstruction, we use metrics *Recall*, *Precision*, *Predicted Proportion*, and *Reconstruction Rate*. *Recall* is defined as the ratio of the number of correctly reconstructed strains to the total number of true strains in the quasispecies, i.e., 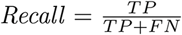, while *Precision* is defined as the fraction of correctly reconstructed strains among all the assembled sequences, i.e., 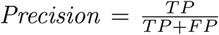. Noting that *Precision* usually reports high scores when the number of strains is underestimated while penalizing overestimation of the population size, we also report the ratio of the number of reconstructed sequences to the true population size, *Predicted Proportion*. The closer *Predicted Proportion* to 1, the more accurate the number of reconstructed strains. Moreover, to assess the degree of reconstruction accuracy, we define 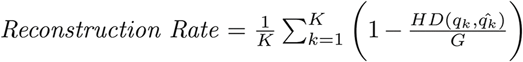, where *G* is the length of a genome, *K* is the number of strains in a quasispecies and *q_k_* and *qˆ*_*k*_ denote the *k^th^* true strain and its nearest sequence among the *K* estimated ones, respectively. To assess the accuracy of estimated frequencies, we use Jensen-Shannon divergence (JSD) which quantifies similarity between two distributions. Given a true distribution *P* and its approximation *Q*, the Kullback-Leibler (KL) divergence 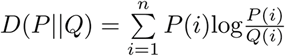 is undefined when *Q*(*i*) = 0. JSD, a symmetrized and smoothed version of the KL divergence, circumvents this problem by defining similarity of *P* and *Q* as 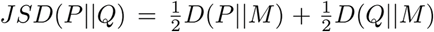, where *M* is defined as 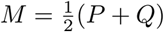.

#### Algorithm 2 Sequential Bayesian Inference for quasispecies reconstruction

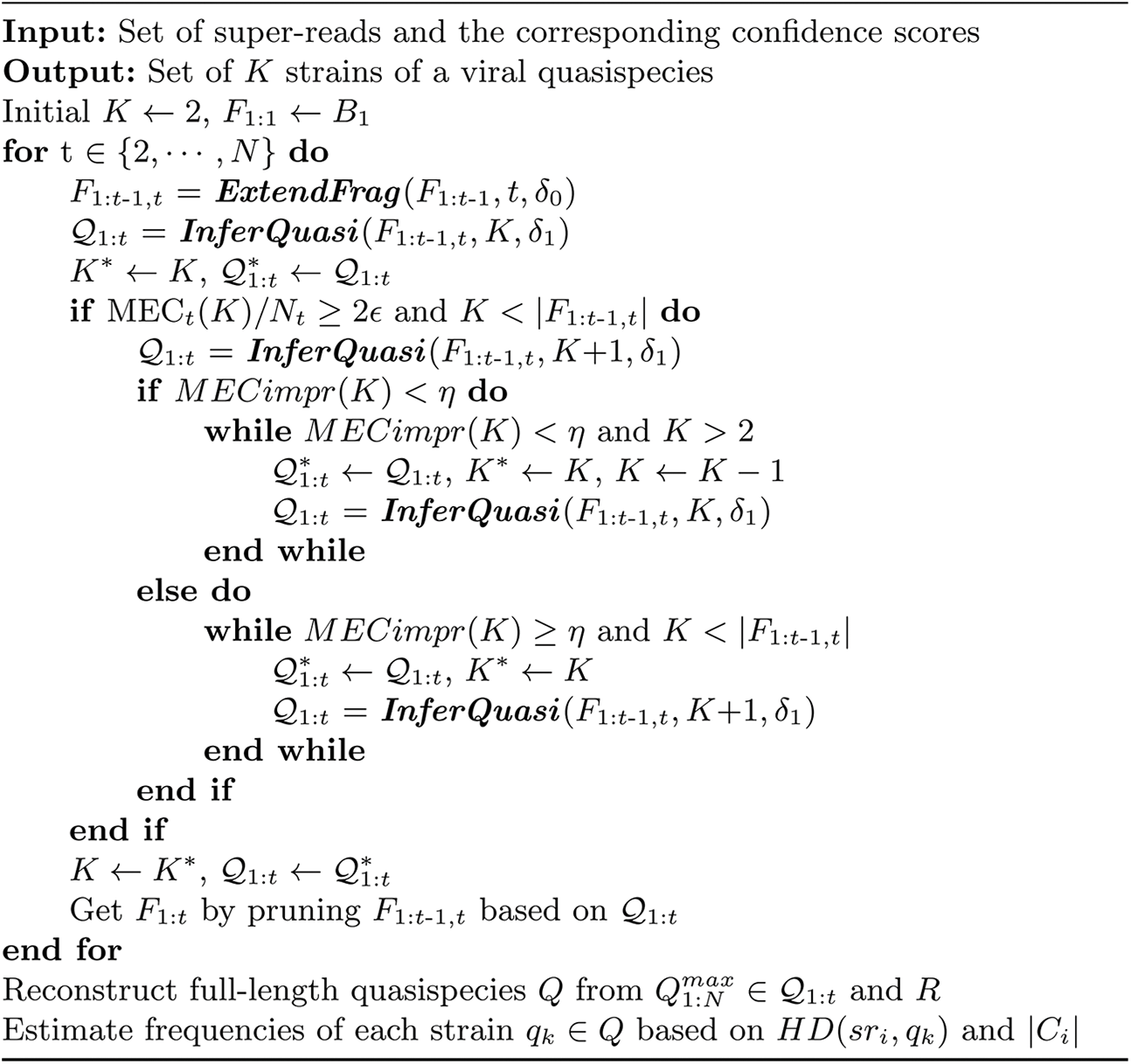

We compare our algorithm with publicly available ShoRAH [4], PredictHaplo [12], and ViQuaS [17]. Since ViQuaS is an extension of the algorithm in [10, 11], and was shown to have superior performance compared to its predecessor, we omit the comparison with the software QuRe in [10, 11]. It is worth pointing out that for the synthetic data sets we study, ShoRAH could not reconstruct strains in the regions where the simulated sequencing coverage is relatively low compared to the average, resulting in reconstruction of strains that are shorter than the true length *G*. To facilitate a fair comparison with ShoRAH, we aligned its reconstructed strains to the reference genome and completed missing sites with bases from the reference. ViQuaS, on the other hand, tends to reconstruct many more strains than actually present; thus we followed ViQuaS’s authors recommendation and retained only those having frequencies greater than *f_min_* when calculating *Precision*. Finally, not all of the synthetic data sets could be processed with PredictHaplo, preventing us from reporting its performance in some of the scenarios.

**Table 1.**
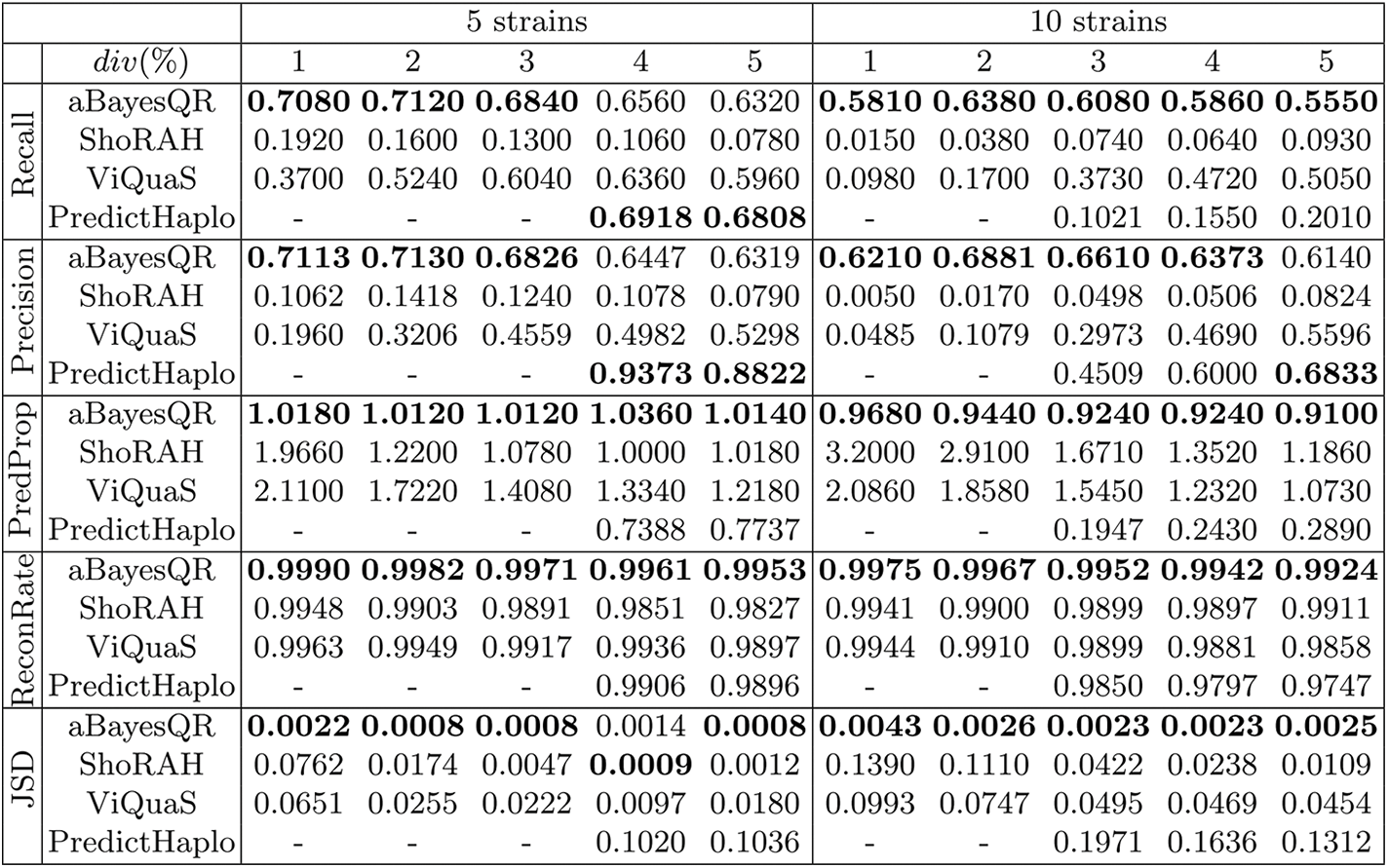
Performance comparison of different methods for varied diversities (*div*) on simulated data. Performance comparison of aBayesQR, ShoRAH, ViQuaS and PredictHaplo in terms of *Recall*, *Precision*, *Predicted Proportion* (*PredProp*), *Reconstruction Rate* (*ReconRate*) and *JSD* on the simulated data with *err* = 0.1% and *cov* = 500 × vs. *div* for a mixture of 5 and 10 viral strains. Averaged PredictHaplo results are reported if it provides answers for more than 50% of data sets. Boldface values indicate the best performance for each *div*(%).

We generated synthetic datasets by emulating high-throughput sequencing of a viral population consisting of a number of closely related viral genomes having length of 1300bp; this particular length was chosen to coincide with the longest region of the HIV *pol* gene. Quasispecies sequences are generated by introducing independent mutations at uniformly random locations along the length of a randomly generated reference genome so as to obtain a predefined level of diversity (*div*%), i.e., a predefined average Hamming distance between quasispecies strains. Simulating Illumina’s MiSeq data, 2 × 250bp-long pairedend reads are sampled uniformly from each viral strain with a mean coverage of *cov×* per strain. Inserts of the paired-end reads are on average 150bp long with standard deviation of 30. In our benchmarking tests, we focus on exploring the effects of diversity (*div*%) on the accuracy of the quasispecies reconstruction. Two sets of viral populations are considered: (1) a mix of 5 viral strains with abundance levels 50%, 30%, 15%, 4% and 1%; and (2) a mix of 10 strains with abundance levels 36%, 24%, 16%, 8%, 5.5%, 4%, 3%, 2%, 1% and 0.5%. Note that the abundances are chosen to approximately follow geometric distribution and that the populations include low abundant strains. For each combination of the parameters, 100 data sets were generated and the reported results were obtained by averaging over those data instances. For PredictHaplo, which did not produce results in each instance, the averaged results are reported if more than 50 instances were successfully processed.

In all of the following experiments, potential SNVs are called if their abundance is higher than 1%, which is set relatively high to avoid false positives (FPs); FPs prevent reads to be merged with existing clusters in Sect. 2.1. We execute the function ***ExtendFrag*** with parameter *δ*_0_ = 0.1. Parameter *δ*_1_ in function ***InferQuasi*** is initially set to 0.001, but adaptively increases if the number of combinations of partially reconstructed strains exceeds 10000; this is done to limit the number of likelihood calculations performed in each run of ***InferQuasi***. The recommended value of *η*, a threshold used to determine population size *K* based on *MECimpr*(·), is discussed in Appendix B.

We compare performances of aBayesQR, ShoRAH, ViQuaS and PredictHap when applied to the reconstruction of a quasispecies spectrum with diversity levels varying between 1% and 5% (i.e., *div* ∈ 1%, 2%, 3%, 4%, 5%). To test the ability of different methods to reconstruct quasispecies with low diversity, we assume low sequencing error rate of *err* = 0.1% (median mismatch error rates for 454 Life Sciences and Illumina platforms are 0.1% and 0.12%, respectively ([25])). Coverage per strain *cov* × is set to 500x, implying total coverage of 2500 and 5000 for the 5-strain and 10-strain population, respectively; strains having frequencies 0.23% or higher in the 5-strain case and those with frequencies 0.46% or higher in the 10-strain case are covered with probability 0.99 ([5]).

Table 1 demonstrates that the proposed aBayesQR algorithm outperforms existing schemes. In terms of *Recall* and *Precision*, aBayesQR exhibits exceptionally good performance compared to competing methods when reconstructing quasispecies strains with diversity *div* < 4%. The performance of ViQuaS deteriorates at low diversities in terms of most of the criteria (i.e., *Recall*, *Precision*, *Predicted Proportion* and *JSD*). PredictHaplo could not perform reconstruction in most of the low diversity instances yet it overall achieves the highest *Precision* because it typically underestimates the number of strains as shown by *Predicted Proportion* (e.g., estimating only 2-3 out of 10 strains), which is in agreement with the results reported by a previous study [14]. Among all methods, ShoRAH has the lowest performance in terms of *Recall* and *Precision*. As indicated by *Predicted Proportion*, aBayesQR is the most accurate method in terms of estimating the population size although it often misses a strain with the lowest frequency when applied to reconstruction of a quasispecies consisting of 10 strains. ViQuaS and ShoRAH typically overestimates the number of strains especially at low diversity levels. aBayesQR is the best method in terms of *Reconstruction Rate* at all levels of diversity. In terms of frequency estimation, aBayesQR overall outperforms all the other methods whereas PredictHaplo shows the highest *JSD* due to its drawback of underestimating the number of strains. Note that both ViQuaS and ShoRAH exhibit significantly increased (i.e., deteriorated) *JSD* at low diversity levels. This fact, along with the low *Recall* and *Precision* scores they have in low diversity settings, indicates that state-of-the-art methods experience major difficulties when attempting to reconstruct viral quasispecies in those settings, as also observed in [5, 14] and [17].

We further study the effects that sequencing error rate (*err*%) and coverage per strain (*cov* ×) have on the performance of the algorithms. Those results are reported in Table S2 and S3 in the Appendix C, demonstrating superiority of aBayesQR as compared to the competing methods. The runtimes of the tested algorithms are shown in Table S4 in the Appendix C.

### 3.2 Performance comparison on real HIV data

To further test the performance of our proposed method, we employ it for the analysis of the HIV 5-virus-mix dataset published in [20]. Specifically, we apply our algorithm to reconstruct an *in vitro* generated quasispecies population consisting of 5 known HIV-1 strains: HIV-1_*HXB*2_, HIV-1_89.6_, HIV-1*_JR−CSF_*, HIV-1_*NL*4−3_ and HIV-1_*YU*2_. Compared to the simulated data set, relative frequencies of the 5 HIV-1 strains are more evenly distributed (about 10% − 30%) and the pairwise distances between strains are higher (2.61% − 8.45%) [20]. We use the 2 × 250bp-long paired-end reads provided by Illumina’s MiSeq Benchtop Sequencer. The reads are aligned to the HIV-1_*HXB*2_ reference genome; the reads shorter than 150nt and those having bases with quality scores less than a PHRED threshold of 60 are discarded. We compare the performance of our method applied to gene-wise quasispecies reconstruction of the above described HIV data with that of the competing techniques. Since the current version of ViQuaS software does not support specifying genomic regions, we could not use it in this experiment. When running aBayesQR, we set the parameter *η* to 0.09 (the setting recommended in Appendix B). Other parameters are set to the same values as the ones used in Sect. 3.1.

We evaluate and report the *Predicted Proportion* (i.e., the fraction of correctly estimated strains as defined in Sect. 3.1) and *Reconstruction Rate* in Table 2. On this real HIV-1 data set which (as pointed above) has different properties than the simulated data set in Sect. 3.1, aBayesQR is the most accurate among the considered methods in terms of *Predicted Proportion*. PredictHaplo underestimates the population size and reconstructs three or four strains in the 8 considered genes and ShoRAH greatly overestimates the population size for all 13 genes of the HIV-1 data set (e.g., it reconstructs 119 strains in gp120), which is consistent with our simulation results as well as with the results in [14]. aBayesQR and PredictHaplo are tied for the number of genes where all the strains are perfectly reconstructed (5 each); for the remaining genes, PredictHaplo provides a larger number of perfectly reconstructed strains. However, it is worth pointing out that PredictHaplo, designed for identification of HIV haplotypes, missed at least one strain in each of the remaining 8 genes while aBayesQR reconstructed most of the strains on all but two genes, gp120 and gp41. ShoRAH did not perfectly reconstruct any of the 13 genes, which is consistent with the simulation results. Moreover, overestimating the number of strains negatively affects the accuracy of ShoRAH’s frequency estimation; for instance, the sum of frequencies corresponding to the most abundant 5 strains does not exceed 50% in 9 out 13 genes (71% is the largest such sum, on *vpu*) (see Table S5 in the Appendix C).

**Table 2.** Performance comparisons on a real HIV-1 5-virus-mix data set. *Predicted Proportion* (PredProp) and *Reconstruction Rate* (RR) for aBayesQR, ShoRAH and PredictHaplo applied to reconstruction of HIV-1_HXB2_, HIV-1_89.6_, HIV-1_JR-CSF_, HIV-1_NL4-3_ and HIV-1_YU2_ for all 13 genes of the HIV-1 dataset. (note: RR are expressed in percentages.) Boldface values indicate the genes where all the strains are perfectly reconstructed. The inferred frequencies are shown in Table S5 in Appendix C.

|  |  | p17 | p24 | p2-p6 | PR | RT | RNase | int | vif | vpr | vpu | gp120 | gp41 | nef |
| --- | --- | --- | --- | --- | --- | --- | --- | --- | --- | --- | --- | --- | --- | --- |
| aBayesQR | PredProp | **1** | **1** | **1** | **1** | **1** | **1** | **1** | **1** | **1.2** | **1** | 0.8 | 0.8 | 1.2 |
|  | RR<sub>HXB2</sub> | **100** | 99.4 | **100** | **100** | 98.5 | **100** | 99.9 | 100 | **100** | 99.6 | 98 | 0 | 95.8 |
|  | RR<sub>89.6</sub> | **100** | 98.7 | **100** | **100** | 98.6 | **100** | 100 | 100 | **100** | 92 | 96.5 | 98.9 | 95.5 |
|  | RR<sub>JR-CSF</sub> | **100** | 99.6 | **100** | **100** | 99 | **100** | 100 | 100 | **100** | 98.8 | 97.7 | 99.1 | 98.2 |
|  | RR<sub>NL4-3</sub> | **100** | 100 | **100** | **100** | 98.9 | **100** | 100 | 99.8 | **100** | 100 | 96.3 | 98.8 | 100 |
|  | RR<sub>YU2</sub> | **100** | 99.7 | **100** | **100** | 99.2 | **100** | 99.5 | 99.7 | **100** | 100 | 0 | 98.6 | 99.2 |
| ShoRAH | PredProp | 13 | 16.4 | 13.8 | 8.8 | 21.8 | 11.8 | 13.6 | 12.8 | 7.8 | 4 | 23.8 | 19.8 | 17.4 |
|  | RR<sub>HXB2</sub> | 100 | 96.7 | 100 | 100 | 98.2 | 100 | 97.5 | 100 | 100 | 100 | 97.7 | 98.4 | 98.2 |
|  | RR<sub>89.6</sub> | 100 | 99.7 | 100 | 100 | 98.6 | 100 | 98.9 | 99.8 | 100 | 93.6 | 96.1 | 98.6 | 98.9 |
|  | RR<sub>JR-CSF</sub> | 100 | 100 | 100 | 100 | 99.8 | 96.4 | 99 | 100 | 100 | 98 | 96.9 | 96.3 | 94.7 |
|  | RR<sub>NL4-3</sub> | 100 | 99.1 | 97.3 | 100 | 98.9 | 99.2 | 99.3 | 99.3 | 100 | 100 | 96.1 | 98.5 | 98.6 |
|  | RR<sub>YU2</sub> | 94.2 | 99 | 100 | 98.3 | 98 | 94.5 | 98.6 | 95 | 93.2 | 90.8 | 97 | 95.4 | 97.9 |
| PredictHaplo | PredProp | **1** | 0.6 | **1** | **1** | **1** | 0.8 | 0.8 | 0.8 | **1** | 0.8 | 0.8 | 0.8 | 0.8 |
|  | RR<sub>HXB2</sub> | **100** | 0 | **100** | **100** | **100** | 98.9 | 100 | 100 | **100** | 93.17 | 0 | 0 | 0 |
|  | RR<sub>89.6</sub> | **100** | 100 | **100** | **100** | **100** | 100 | 99.8 | 100 | **100** | 0 | 97.8 | 100 | 98.87 |
|  | RR<sub>JR-CSF</sub> | **100** | 100 | **100** | **100** | **100** | 100 | 100 | 100 | **100** | 100 | 99.7 | 100 | 100 |
|  | RR<sub>NL4-3</sub> | **100** | 99.1 | **100** | **100** | **100** | 100 | 100 | 100 | **100** | 100 | 100 | 100 | 100 |
|  | RR<sub>YU2</sub> | **100** | 0 | **100** | **100** | **100** | 0 | 0 | 0 | **100** | 100 | 98.6 | 100 | 100 |

To complement the gene-wise quasispecies reconstruction study with that of a global reconstruction, we consider the HIV-1 *gap-pol* region spanning 4307bp. To efficiently process 355241 paired-end reads that remain after applying a quality filter, we organize the region into a sequence of windows of length 400bp where the consecutive windows overlap by 150bp and run aBayesQR on those windows. The entire region is assembled by connecting strains in the consecutive windows while testing consistency in the overlapping intervals. The number of strains retrieved in the global reconstruction is decided by majority voting of the number of strains obtained in each window. The frequencies are estimated by counting reads nearest (in terms of Hamming distance) to each of the reconstructed strains. Following this procedure, both aBayesQR and PredictHaplo could reconstruct all 5 HIV-1 strains in the *gap-pol* region correctly, i.e., they both achieved *Reconstruction rate* of 100 for all 5 strains and *Predicted Proportion* of 1. The frequencies estimated by aBayesQR are 15.21%, 19.34%, 25.56%, 27.61% and 12.27% while those estimated by PredictHap are 13.21%, 13.56%, 25.67%, 19.69% and 27.86%. ShoRAH highly overestimated the number of strains and reported *Predicted Proportion* of 41.8; its five most abundant strains estimated are reported to have frequencies 8.51%, 5.04%, 3.41%, 3.24% and 3.09%.

## 4. Conclusions

In this paper, we presented a novel maximum-likelihood based approximate algorithm for reconstructing viral quasispecies from high-throughput sequencing data. aBayesQR assembles paired-end short reads into longer fragments based on similarity of the read overlaps and the uncertainty level of non-overlapping regions. The probable sets of partially reconstructed strains are inductively searched and a subset of those strains is extended to efficiently deduce the most likely set of strains in a quasisepcies. Detection of the population size is embedded into the algorithm and is empirically shown to be very accurate; the number of strains is dynamically adjusted based on the reliability of the partially assembled quasispecies in each extension step. Performance of the developed method is tested on both synthetic datasets and a real HIV-1 dataset. In both settings, the new algorithm outperforms existing techniques in terms of accuracy of the quasispecies size estimation, perfect reconstruction of strains, proportion of correct bases in each reconstructed strain and the estimation of their abundance. A particularly high accuracy is observed in estimating the population size (i.e., the number of strains) and their relative abundance. Tests on synthetic datasets demonstrates that aBayesQR is capable of reconstructing quasispecies at low diversity, showing superior performance in those settings compared to state-of-the-art algorithms. Furthermore, the study on a real HIV-1 dataset demonstrates that our proposed algorithm outperforms or has performance comparable to that of the existing methods in the general setting of viral quasispecies reconstruction.

aBayesQR can be extended and applied to the problem of estimating the population size and the degree of variation among the constituent species in related fields such as immunogenetics. On a related note, bacterial populations are characterized by having relatively lower mutation rates than viral and thus typically have fewer segregating sites on the sequences in a population. The ability of our method to perform highly accurate reconstruction in such settings should be further investigated.

A software aBayesQR is available at https://github.com/SoyeonA/aBayesQR.

### Acknowledgements

This work was funded by the National Science Foundation under grants CCF 1507998 and CCF 1618427.

## Footnotes

1 If either of the two sequences has a gap “*−*” in a position, that position is ignored in the computation of the aforementioned Hamming distance.

2 This matches typical variations in the HIV *pol* gene which range between 3% and 5% ([5]).

## Appendix A

**Fig. S1.**
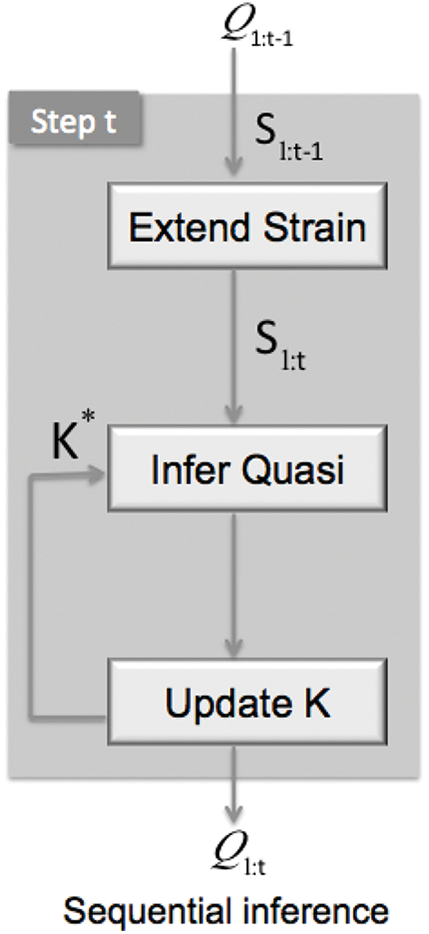
Procedure of sequential Bayesian inference in step *t*

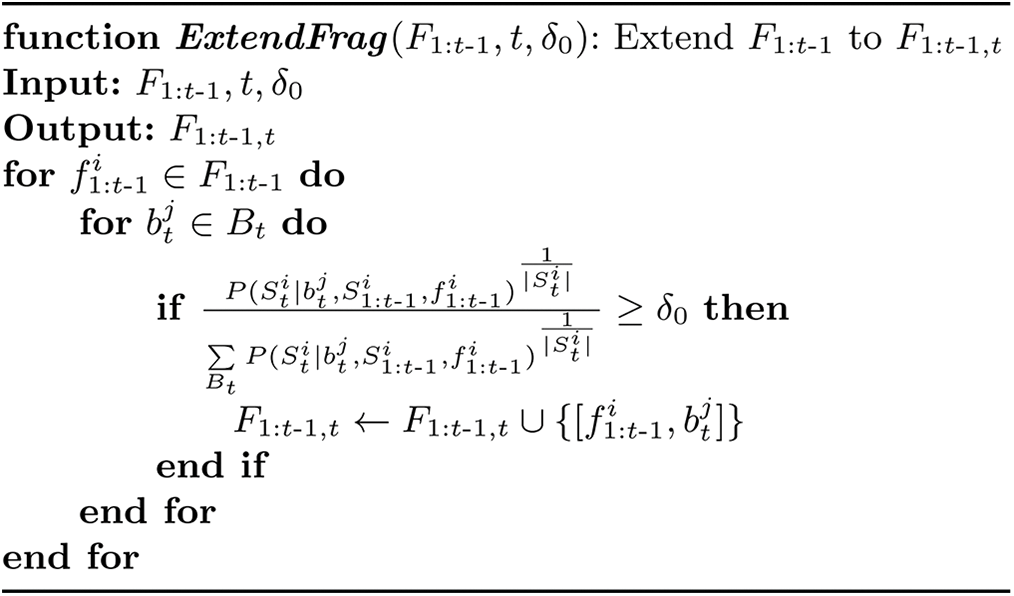

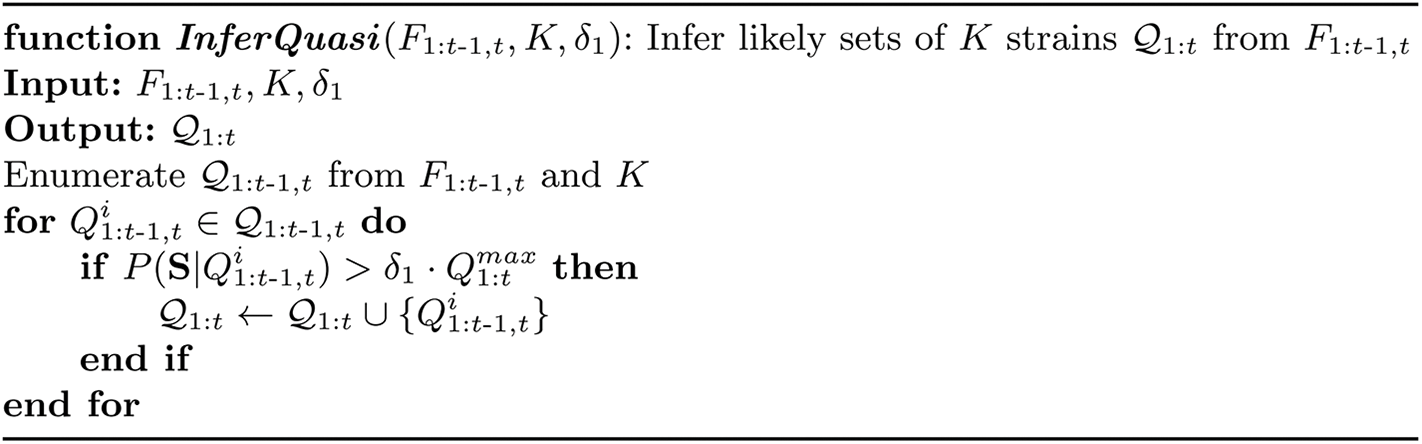

## Appendix B: Guideline for choosing parameter *η*

Here we discuss the choice of parameter *η*, a threshold used for assessing value of the metric *MECimpr* in the process of determining the number of sequences *K* in a quasispecies. Let *Q* = {*q_k_*, *k* = 1,…, *K*} denote a viral population consisting of K strains. The set of reads of length *L, R* = {*r_i_*, *i* = 1,…, |*R*|}, is generated by a sequencing platform having error rate *є* and aligned to the reference genome of length *G*. The MEC score characterizing accuracy of the assembly of strains in *Q* from reads in *R* is calculated as

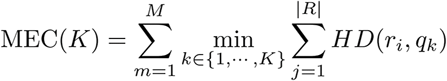

Assume that the sequencing errors are independent and identically distributed across all reads. When the quasispecies recovery is perfect, i.e., when all of the *K* strains are reconstructed correctly and the relative frequencies of strains are estimated accurately, the MEC score is *|R|L*ε. If a strain with the smallest frequency (*f_min_*) is not recognized as a distinct mixture component while the other *K* − 1 sequences are correctly reconstructed, the incorrect clustering of |*R| f_min_* reads generated from the rarest quasispecies component induces an extra contribution to the MEC score. The MEC score obtained following perfect reconstruction of *K* − 1 strains and misclassification of the *K^th^* strain is

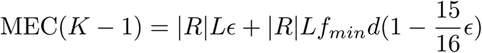

where *d* is the average diversity rate. The MEC improvement rate achieved by increasing the number of viral strains from *K* – 1 to *K* (and thus reducing the MEC score thanks to a correct clustering of the rarest strain) is

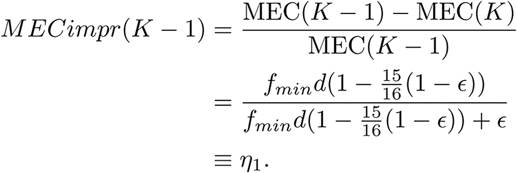

While *η*_1_ is a potential choice for the threshold, it is beneficial to soften it and allow that *MECimpr*(·) takes on values slightly below it. To this end, let us also consider the scenario where in addition to the perfect recovery of *K* strains, an extra strain is erroneously inferred by misclassifying reads that in fact should have been placed in the cluster associated with the most abundant strain (i.e., the one having frequency *f_max_*). This reduces evaluated MEC score to an unrealistically low value given by

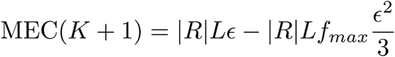

Improvement of the MEC score due to having an extra (unnecessary) cluster can be expressed as

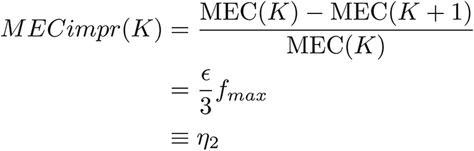

We note that for typical parameter values, *η*1 » *η*2; we choose the threshold *η* by taking a weighted geometric mean of *η*_1_ and *η*_2_,

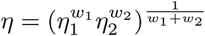

To avoid overestimation of the number of strains, we choose *w*_1_ to be larger than *w*_2_. In our experiments, the ratio *r* = *w*_1_*/w*_2_ was set to 5. We find that the results are fairly robust with respect to the choice of parameter *η* as demonstrated in Table S1.

**Table S1.** Performances comparison of aBayesQR with different parameter *η* for varied diversities *div* on simulated data. Performances of aBayesQR as a function of diversity with parameter *η* varied around the recommended value from −20% to +20%; shown are *Recall*, *Precision*, *Predicted Proportion* (*PredProp*), *Reconstruction Rate* (*Recon-Rate*) and *JSD*. The data is generated synthetically, relevant parameters are *err* = 0.1% and *cov* = 500×, simulated is a mixture of 5 and 10 viral strains.

|  |  | 5 strains |  |  |  |  | 10 strains |  |  |  |  |
| --- | --- | --- | --- | --- | --- | --- | --- | --- | --- | --- | --- |
| | $div(\%)$ | 1 | 2 | 3 | 4 | 5 | 1 | 2 | 3 | 4 | 5 |
| Recall | $0.8\eta$ | 0.7020 | 0.7080 | 0.6840 | 0.6560 | 0.6320 | 0.5800 | 0.6390 | 0.6060 | 0.5810 | 0.5550 |
| | $0.9\eta$ | 0.7060 | 0.7120 | 0.6840 | 0.6560 | 0.6300 | 0.5800 | 0.6390 | 0.6080 | 0.5850 | 0.5550 |
| | $\eta$ | 0.7080 | 0.7120 | 0.6840 | 0.6560 | 0.6320 | 0.5810 | 0.6380 | 0.6080 | 0.5860 | 0.5550 |
| | $1.1\eta$ | 0.7080 | 0.7120 | 0.6840 | 0.6560 | 0.6300 | 0.5780 | 0.6390 | 0.6100 | 0.5850 | 0.5580 |
| | $1.2\eta$ | 0.7060 | 0.7120 | 0.6840 | 0.6560 | 0.6340 | 0.5780 | 0.6370 | 0.6100 | 0.5850 | 0.5590 |
| Precision | $0.8\eta$ | 0.7019 | 0.7069 | 0.6811 | 0.6397 | 0.6265 | 0.6186 | 0.6874 | 0.6553 | 0.6273 | 0.6094 |
| | $0.9\eta$ | 0.7083 | 0.7130 | 0.6811 | 0.6427 | 0.6301 | 0.6190 | 0.6892 | 0.6602 | 0.6325 | 0.6110 |
| | $\eta$ | 0.7113 | 0.7130 | 0.6826 | 0.6447 | 0.6319 | 0.6210 | 0.6881 | 0.6610 | 0.6373 | 0.6140 |
| | $1.1\eta$ | 0.7113 | 0.7144 | 0.6826 | 0.6470 | 0.6301 | 0.6177 | 0.6892 | 0.6637 | 0.6379 | 0.6238 |
| | $1.2\eta$ | 0.7089 | 0.7144 | 0.6841 | 0.6448 | 0.6410 | 0.6181 | 0.6889 | 0.6660 | 0.6398 | 0.6265 |
| PredProp | $0.8\eta$ | 1.0260 | 1.0160 | 1.0140 | 1.0400 | 1.0240 | 0.9720 | 0.9470 | 0.9300 | 0.9310 | 0.9180 |
| | $0.9\eta$ | 1.0200 | 1.0120 | 1.0140 | 1.0380 | 1.0160 | 0.9710 | 0.9440 | 0.9250 | 0.9290 | 0.9140 |
| | $\eta$ | 1.0180 | 1.0120 | 1.0120 | 1.0360 | 1.0140 | 0.9680 | 0.9440 | 0.9240 | 0.9240 | 0.9100 |
| | $1.1\eta$ | 1.0180 | 1.0100 | 1.0120 | 1.0320 | 1.0160 | 0.9690 | 0.9440 | 0.9230 | 0.9210 | 0.9020 |
| | $1.2\eta$ | 1.0200 | 1.0100 | 1.0100 | 1.0340 | 1.0060 | 0.9680 | 0.9410 | 0.9200 | 0.9180 | 0.8990 |
| ReconRate | $0.8\eta$ | 0.9990 | 0.9980 | 0.9971 | 0.9962 | 0.9954 | 0.9975 | 0.9967 | 0.9952 | 0.9941 | 0.9924 |
| | $0.9\eta$ | 0.9990 | 0.9982 | 0.9971 | 0.9962 | 0.9953 | 0.9975 | 0.9967 | 0.9952 | 0.9943 | 0.9924 |
| | $\eta$ | 0.9990 | 0.9982 | 0.9971 | 0.9961 | 0.9953 | 0.9975 | 0.9967 | 0.9952 | 0.9942 | 0.9924 |
| | $1.1\eta$ | 0.9990 | 0.9982 | 0.9971 | 0.9961 | 0.9953 | 0.5951 | 0.6621 | 0.6350 | 0.6094 | 0.5879 |
| | $1.2\eta$ | 0.9990 | 0.9982 | 0.9971 | 0.9961 | 0.9951 | 0.9975 | 0.9967 | 0.9952 | 0.9942 | 0.9922 |
| JSD | $0.8\eta$ | 0.0022 | 0.0016 | 0.0009 | 0.0010 | 0.0007 | 0.0043 | 0.0025 | 0.0021 | 0.0021 | 0.0023 |
| | $0.9\eta$ | 0.0022 | 0.0008 | 0.0009 | 0.0014 | 0.0008 | 0.0043 | 0.0026 | 0.0022 | 0.0021 | 0.0024 |
| | $\eta$ | 0.0022 | 0.0008 | 0.0008 | 0.0014 | 0.0008 | 0.0043 | 0.0026 | 0.0023 | 0.0023 | 0.0025 |
| | $1.1\eta$ | 0.0022 | 0.0008 | 0.0008 | 0.0014 | 0.0008 | 0.0043 | 0.0026 | 0.0023 | 0.0023 | 0.0030 |
| | $1.2\eta$ | 0.0022 | 0.0008 | 0.0008 | 0.0015 | 0.0010 | 0.0043 | 0.0026 | 0.0024 | 0.0024 | 0.0030 |

## Appendix C: Supplementary results

**Table S2.**
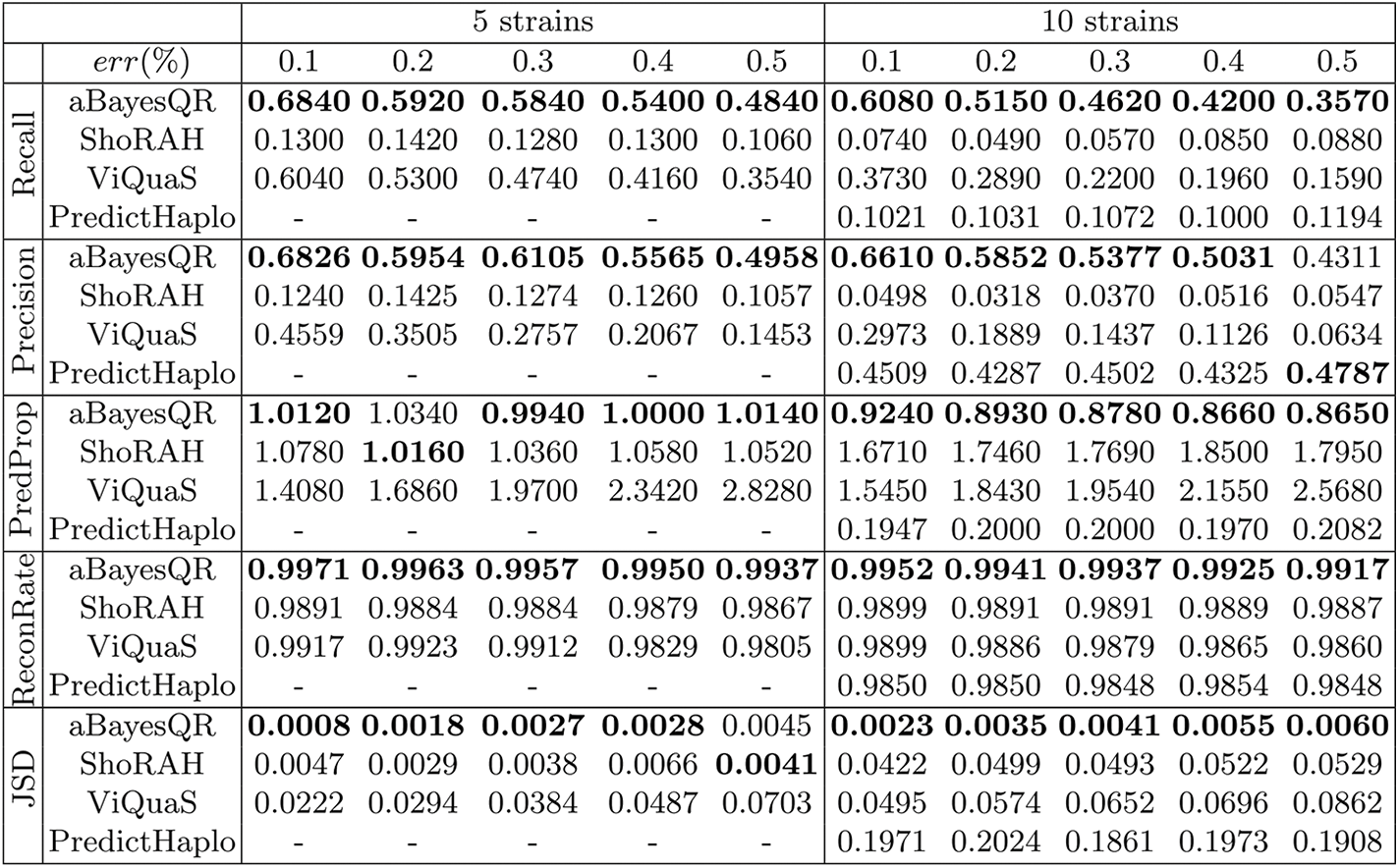
Performance comparison of different methods for varied error rates (*err*) on simulated data. Performance comparison of aBayesQR, ShoRAH, ViQuaS and PredictHaplo in terms of *Recall*, *Precision*, *Predicted Proportion* (*PredProp*), *Reconstruction Rate* (*ReconRate*) and *JSD* on the simulated data with *div* = 3% and *cov* = 500× vs. *err* for a mixture of 5 and 10 viral strains. Averaged PredictHaplo results are reported if it provides answers for more than 50% of data sets. Boldface value indicate the best performance for each *err*(%).

In Table S2, we report the results of the study of the effects sequencing errors have on the performance of aBayesQR, ShoRAH, ViQuaS and PredictHaplo. In particular, the sequencing error is varied from 0.1% to 0.5% (specifically, *err* ∈ {0.1%, 0.2%, 0.3%, 0.4%, 0.5%}), reflecting the range of errors observed in the current and anticipated in future NSG technologies (e.g., the error rates of Illumina’s Miseq have been reported to be below 0.4% in [26] and as high as 0.49 in [27]). We set *div* to be 3%^2^ and set *cov* to be 500× which is the same as in the first experiment in 3.1. In this set of experiments, aBayesQR outperforms ShoRAH, ViQuaS and PredictHaplo over the considered range of *err* achieving the best scores overall for all 5 metrics. As expected, the performances of all methods deteriorate as *err* increases. Since PredictHaplo failed to generate results in most of the instances of the reconstruction problem involving a mixture of 5 strains, its results are not reported. For the problem involving a mixture with 10 strains, PredictHaplo did run successfully in most of the instances but significantly underdetermined the number of strains; on average, its *Predicted Proportion* is around 0.2 (while its *Precision*, as argued earlier in the paper, is somewhat misleadingly good). ViQuaS overestimates the number of strains in all instances; we observe that as *err* increases, ViQuaS generates increasingly more false negative viral strains which adversely affects *Precision* and *JSD*. Even though ShoRAH exhibits the lowest perfect reconstruction scores, it achieves better performance than ViQuaS in terms of frequency estimation and the number of reconstructed strains (i.e., *Predicted Proportion* and *JSD*) when applied to reconstruction of mixtures with 5 strains. For a mixture of 10 strains, however, ShoRAH overestimated the number of strains, which also leads to higher *JSD* scores.

**Table S3.** Performance comparison of different methods for varied coverages (*cov*) on simulated data. Performance comparison of aBayesQR, ShoRAH, ViQuaS and PredictHaplo in terms of *Recall*, *Precision*, *Predicted Proportion* (*PredProp*), *Reconstruction Rate* (*ReconRate*) and *JSD* on the simulated data with *div* = 3% and *err* = 0.3% vs. *cov* for a mixture of 5 and 10 viral strains. Averaged PredictHaplo results are reported if it provides answers for more than 50% of data sets. Boldface value indicate the best performance for each *cov*(×).

|  |  | 5 strains |  |  | 10 strains |  |  |
| --- | --- | --- | --- | --- | --- | --- | --- |
| | *cov*(×) | 250 | 500 | 750 | 250 | 500 | 750 |
| Recall | aBayesQR | **0.4440** | **0.5840** | **0.5920** | **0.4450** | **0.4620** | **0.4450** |
|  | ShoRAH | 0.1820 | 0.1280 | 0.0900 | 0.0590 | 0.0570 | 0.3220 |
|  | ViQuaS | 0.4220 | 0.4740 | 0.4840 | 0.1970 | 0.2200 | 0.2580 |
|  | PredictHaplo | - | - | - | 0.1070 | 0.1072 | 0.1094 |
| Precision | aBayesQR | **0.4231** | **0.6105** | **0.6393** | **0.5131** | **0.5377** | **0.5195** |
|  | ShoRAH | 0.1831 | 0.1274 | 0.0979 | 0.0544 | 0.0370 | 0.2136 |
|  | ViQuaS | 0.2291 | 0.2757 | 0.3256 | 0.0965 | 0.1437 | 0.1900 |
|  | PredictHaplo | - | - | - | 0.4558 | 0.4502 | 0.4635 |
| PredProp | aBayesQR | 1.0920 | **0.9940** | **0.9400** | **0.8840** | **0.8780** | **0.8800** |
|  | ShoRAH | **1.0140** | 1.0360 | 1.0660 | 1.1200 | 1.7690 | 1.6440 |
|  | ViQuaS | 1.9240 | 1.9700 | 1.6820 | 2.2940 | 1.9540 | 1.7180 |
|  | PredictHaplo | - | - | - | 0.2030 | 0.2000 | 0.2031 |
| ReconRate | aBayesQR | **0.9941** | **0.9957** | **0.9959** | **0.9939** | **0.9937** | **0.9930** |
|  | ShoRAH | 0.9906 | 0.9884 | 0.9854 | 0.9874 | 0.9891 | 0.9926 |
|  | ViQuaS | 0.9905 | 0.9912 | 0.9918 | 0.9861 | 0.9879 | 0.9881 |
|  | PredictHaplo | - | - | - | 0.9854 | 0.9848 | 0.9854 |
| JSD | aBayesQR | 0.0040 | **0.0027** | **0.0026** | **0.0041** | **0.0041** | **0.0049** |
|  | ShoRAH | **0.0021** | 0.0038 | 0.0100 | 0.0051 | 0.0493 | 0.0330 |
|  | ViQuaS | 0.0454 | 0.0384 | 0.0318 | 0.0839 | 0.0652 | 0.0555 |
|  | PredictHaplo | - | - | - | 0.1946 | 0.1861 | 0.1930 |

In Table S3, we report the performance of the proposed algorithm for different coverage per strain, *cov* ∈ {250×, 500×, 750×}, while fixing other parameters – specifically, diversity is set to 3%, which is the same as in the second experiment in Table S2, and sequencing error rate is set to 0.3%, which emulates the error rates of Illumina’s Miseq (*<* 0.4% [26]). Performance of four algorithms as a function of coverage is compared in Table S3, demonstrating superiority of aBayesQR in all five metrics of interest.

**Table S4.** Running time comparisons (sec). Running time comparisons of aBayesQR, ShoRAH, ViQuaS and PredictHaplo on the simulated data with *cov* ∈ {250×, 500×, 750×}, *div*=3% and *err*=0.3%, measured on a Linux OS desktop with 3.06GHz CPU and 8Gb RAM (Intel Core i7 880 processor). PredictHaplo results are shown if it provides answers for more than 50% of data sets.

|  | 5 strains |  |  | 10 strains |  |  |
| --- | --- | --- | --- | --- | --- | --- |
| $cov(\times)$ | 250 | 500 | 750 | 250 | 500 | 750 |
| aBayesQR | 96 | 113 | 236 | 451 | 739 | 1606 |
| ShoRAH | 559 | 2005 | 5265 | 2716 | 11473 | 22923 |
| ViQuaS | 93 | 337 | 718 | 360 | 1300 | 13350 |
| PredictHaplo | - | - | - | 93 | 143 | 187 |

Runtimes of each of the algorithms applied to this test set as a function of *cov* are shown in Table S4; note that this characterization of speed (i.e., complexity vs. *cov*) is the most meaningful one to study since the coverage is a main factor affecting the runtime of performing a reconstruction task. The speed is measured on a Linux OS desktop with 3.06GHz CPU and 8GbRAM (Intel Core i7 880 processor). When it completes the task and provides a solution, PredictHaplo is the most efficient among all schemes; however, this method fails to provide answers in most of instances on a mixture of 5 strains. Among the remaining 3 algorithms, our aBayesQR demonstrates the best time efficiency for *cov* ≥ 500 while ShoRAH is the slowest one. ViQuaS is relatively fast at the low coverage *cov* = 250 but its time complexity appears to grow exponentially as *cov* increases, especially in the setting with a mixture of 10 viral strains.

**Table S5.**
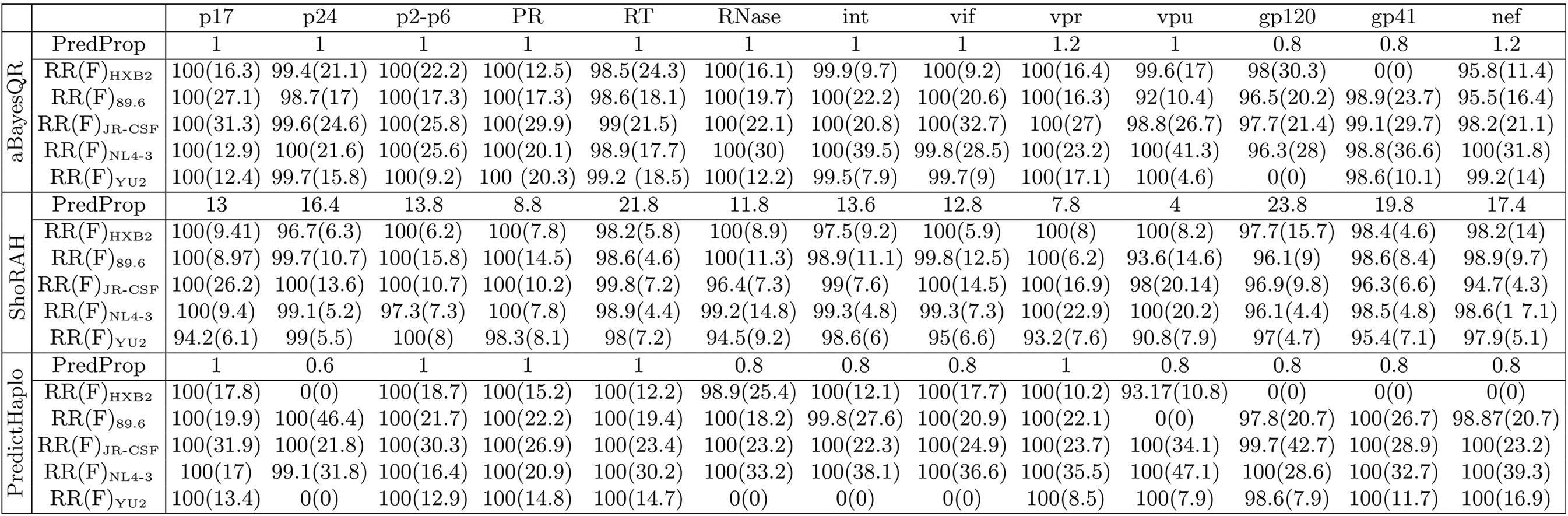
Performance comparisons on a real HIV-1 5-virus-mix data set. *Predicted Proportion* (PredProp), *Reconstruction Rate* (RR) and inferred frequencies (F) for aBayesQR, ShoRAH and PredictHaplo applied to reconstruction of HIV-1_HXB2_, HIV-1_89.6_, HIV-1_JR-CSF_, HIV-1_NL4-3_ and HIV-1_YU2_ for all 13 genes of the HIV-1 dataset (note: frequencies are reported in parenthesis, both RR and F are expressed in percentages).

## References

1 Duarte, E., Novella, I., Weaver, S., Domingo, E., Wain‐Hobson, S., Clarke, D., Moya, A., Elena, S., De La Torre, J., Holland, J.: Rna virus quasispecies: significance for viral disease and epidemiology. Infectious agents and disease 3(4), 201–214 (1994)

2. Lauring, A.S., Andino, R.: Quasispecies theory and the behavior of rna viruses. PLoS Pathogens 6(7) (2010)

3. Posada‐Cespedes, S., Seifert, D., Beerenwinkel, N.: Recent advances in inferring viral diversity from high‐throughput sequencing data. Virus Research (2016)

4. Zagordi, O., Bhattacharya, A., Eriksson, N., Beerenwinkel, N.: Shorah: estimating the genetic diversity of a mixed sample from next‐generation sequencing data. BMC bioinformatics 12(1), 11–9 (2011)

5. Eriksson, N., Pachter, L., Mitsuya, Y., Rhee, S.Y., Wang, C., Gharizadeh, B., Ronaghi, M., Shafer, R.W., Beerenwinkel, N.: Viral population estimation using pyrosequencing. PLoS Comput Biol 4(5), e1000,074 (2008)

6. Zagordi, O., Geyrhofer, L., Roth, V., Beerenwinkel, N.: Deep sequencing of a genetically heterogeneous sample: local haplotype reconstruction and read error correction. Journal of computational biology 17(3), 417–428 (2010)

7. Zagordi, O., Klein, R., Däumer, M., Beerenwinkel, N.: Error correction of next‐generation sequencing data and reliable estimation of hiv quasispecies. Nucleic acids research 38(21), 7400–7409 (2010)

8. Astrovskaya, I., Tork, B., Mangul, S., Westbrooks, K., Măndoiu, I., Balfe, P., Zelikovsky, A.: Inferring viral quasispecies spectra from 454 pyrosequencing reads. BMC bioinformatics 12(6), 1 (2011)

9. Westbrooks, K., Astrovskaya, I., Campo, D., Khudyakov, Y., Berman, P., Zelikovsky, A.: Hcv quasispecies assembly using network flows. In: Bioinformatics Research and Applications, pp. 159–170. Springer (2008)

10. Prosperi, M.C., Prosperi, L., Bruselles, A., Abbate, I., Rozera, G., Vincenti, D., Solmone, M.C., Capobianchi, M.R., Ulivi, G.: Combinatorial analysis and algorithms for quasispecies reconstruction using next‐generation sequencing. BMC bioinformatics 12(1), 1 (2011)

11. Prosperi, M.C., Salemi, M.: Qure: software for viral quasispecies reconstruction from next‐generation sequencing data. Bioinformatics 28(1), 132–133 (2012)

12. Prabhakaran, S., Rey, M., Zagordi, O., Beerenwinkel, N., Roth, V.: Hiv haplotype inference using a propagating dirichlet process mixture model. IEEE/ACM Trans. on Comput. Biol. Bioinform. (TCBB) 11(1), 182–191 (2014)

13. Topfer, A., Zagordi, O., Prabhakaran, S., Roth, V., Halperin, E., Beerenwinkel, N.: Probabilistic inference of viral quasispecies subject to recombination. Journal of Computational Biology 20(2), 113–123 (2013)

14. Schirmer, M., Sloan, W.T., Quince, C.: Benchmarking of viral haplotype reconstruction programmes: an overview of the capacities and limitations of currently available programmes. Briefings in bioinformatics p. bbs081 (2012)

15. Topfer, A., Marschall, T., Bull, R.A., Luciani, F., Schönhuth, A., Beerenwinkel, N.: Viral quasispecies assembly via maximal clique enumeration. PLoS Comput Biol 10(3), e1003,515 (2014)

16. Mangul, S., Wu, N.C., Mancuso, N., Zelikovsky, A., Sun, R., Eskin, E.: Accurate viral population assembly from ultra‐deep sequencing data. Bioinformatics 30(12), i329–i337 (2014)

17. Jayasundara, D., Saeed, I., Maheswararajah, S., Chang, B., Tang, S.L., Halgamuge, S.K.: Viquas: an improved reconstruction pipeline for viral quasispecies spectra generated by next‐generation sequencing. Bioinformatics p. btu754 (2014)

18. Le, T., Chiarella, J., Simen, B.B., Hanczaruk, B., Egholm, M., Landry, M.L., Dieckhaus, K., Rosen, M.I., Kozal, M.J.: Low‐abundance hiv drug‐resistant viral variants in treatment‐experienced persons correlate with historical antiretroviral use. PloS one 4(6), e6079 (2009)

19. Simen, B.B., Simons, J.F., Hullsiek, K.H., Novak, R.M., MacArthur, R.D., Baxter, J.D., Huang, C., Lubeski, C., Turenchalk, G.S., Braverman, M.S., et al.: Low‐abundance drug‐resistant viral variants in chronically hiv‐infected, antiretroviral treatment‐naive patients significantly impact treatment outcomes. Journal of Infectious Diseases 199(5), 693–701 (2009)

20. Di Giallonardo, F., Töpfer, A., Rey, M., Prabhakaran, S., Duport, Y., Leemann, C., Schmutz, S., Campbell, N.K., Joos, B., Lecca, M.R., et al.: Full‐length haplotype reconstruction to infer the structure of heterogeneous virus populations. Nucleic acids research 42(14), e115–e115 (2014)

21. Sasirekha, K., Baby, P.: Agglomerative hierarchical clustering algorithm‐a review. International Journal of Scientific and Research Publications 3(3) (2013)

22. Jung, S.Y., Kim, T.S.: An agglomerative hierarchical clustering using partial maximum array and incremental similarity computation method. In: Data Mining, 2001. ICDM 2001, Proc. IEEE Int. Conf. on, pp. 265–272. IEEE (2001)

23. Lancia, G., Bafna, V., Istrail, S., Lippert, R., Schwartz, R.: Snps problems, complexity, and algorithms. In: Algorithms—ESA 2001, pp. 182–193. Springer (2001)

24. Lippert, R., Schwartz, R., Lancia, G., Istrail, S.: Algorithmic strategies for the single nucleotide polymorphism haplotype assembly problem. Briefings in bioinformatics 3(1), 23–31 (2002)

25. Archer, J., Baillie, G., Watson, S.J., Kellam, P., Rambaut, A., Robertson, D.L.: Analysis of high‐depth sequence data for studying viral diversity: a comparison of next generation sequencing platforms using segminator ii. BMC bioinformatics 13(1), 4–7 (2012)

26. Quail, M.A., Smith, M., Coupland, P., Otto, T.D., Harris, S.R., Connor, T.R., Bertoni, A., Swerdlow, H.P., Gu, Y.: A tale of three next generation sequencing platforms: comparison of ion torrent, pacific biosciences and illumina miseq sequencers. BMC genomics 13(1), 1 (2012)

27. Ross, M.G., Russ, C., Costello, M., Hollinger, A., Lennon, N.J., Hegarty, R., Nusbaum, C., Jaffe, D.B.: Characterizing and measuring bias in sequence data. Genome Biol 14(5), R51 (2013)

